# Inhibition of LRRK2 kinase activity promotes anterograde axonal transport and presynaptic targeting of α-synuclein

**DOI:** 10.1101/2021.10.04.463043

**Authors:** Charlotte F. Brzozowski, Baraa A. Hijaz, Vijay Singh, Nolwazi Z. Gcwensa, Kaela Kelly, Edward S. Boyden, Andrew B. West, Deblina Sarkar, Laura A. Volpicelli-Daley

## Abstract

Pathologic inclusions composed of α-synuclein called Lewy pathology are hallmarks of Parkinson’s Disease (PD). Dominant inherited mutations in leucine rich repeat kinase 2 (LRRK2) are the most common genetic cause of PD. Lewy pathology is found in the majority of individuals with LRRK2-PD, particularly those with the G2019S-LRRK2 mutation. Lewy pathology in LRRK2-PD associates with increased non-motor symptoms such as cognitive deficits, anxiety, and orthostatic hypotension. Thus, understanding the relationship between LRRK2 and α-synuclein could be important for determining the mechanisms of non-motor symptoms. In PD models, expression of mutant LRRK2 reduces membrane localization of α- synuclein, and enhances formation of pathologic α-synuclein, particularly when synaptic activity is increased. α-Synuclein and LRRK2 both localize to the presynaptic terminal. LRRK2 plays a role in membrane traffic, including axonal transport, and therefore may influence α-synuclein synaptic localization. This study shows that LRRK2 kinase activity influences α-synuclein targeting to the presynaptic terminal. We used the selective LRRK2 kinase inhibitors, MLi-2 and PF-06685360 (PF-360) to determine the impact of reduced LRRK2 kinase activity on presynaptic localization of α-synuclein. Expansion microscopy (ExM) in primary hippocampal cultures and the mouse striatum, *in vivo*, was used to more precisely resolve the presynaptic localization of α-synuclein. Live imaging of axonal transport of α-synuclein-GFP was used to investigate the impact of LRRK2 kinase inhibition on α-synuclein axonal transport towards the presynaptic terminal. Reduced LRRK2 kinase activity increases α-synuclein overlap with presynaptic markers in primary neurons, and increases anterograde axonal transport of α- synuclein-GFP. *In vivo*, LRRK2 inhibition increases α-synuclein overlap with glutamatergic, cortico-striatal terminals, and dopaminergic nigral-striatal presynaptic terminals. The findings suggest that LRRK2 kinase activity plays a role in axonal transport, and presynaptic targeting of α-synuclein. These data provide potential mechanisms by which LRRK2-mediated perturbations of α-synuclein localization could cause pathology in both LRRK2-PD, and idiopathic PD.

## Introduction

Inclusions composed primarily of the protein, □-synuclein, found in axons, called Lewy neurites, and in the soma, called Lewy bodies [1], are among the primary hallmarks of PD. Mutations in LRRK2 cause familial PD, and increase its kinase activity. Gene variations in LRRK2 increase risk of developing sporadic PD [2, 3]. Increased LRRK2 kinase activity has also been observed in post mortem tissue analysis of idiopathic PD brains [4]. Lewy pathology is common in individuals with PD harboring the most common G2019S-LRRK2 mutation, which is associated with increased nonmotor symptoms [5]. PD animal models demonstrate α-synuclein pathologic aggregation is exacerbated by G2019S-LRRK2 expression [6-11]. Given the importance of LRRK2 activity in both familial, and idiopathic PD cases, deciphering the downstream effects of LRRK2 kinase activity is crucial for investigating PD mechanisms, in general. An understanding of how LRRK2 kinase activity interacts with α-synuclein could help with determine how these two proteins contribute to LRRK2-PD, and idiopathic PD.

α-Synuclein is enriched in presynaptic terminals of glutamatergic neurons, and stabilizes synaptic vesicle SNARE formation via interactions with VAMP2, acting as a brake on synaptic vesicle exocytosis [12-17]. LRRK2 also plays a role in presynaptic vesicle traffic, and functionally interacts with presynaptic vesicle proteins [18-24]. One of the most well-established roles for LRRK2 is in membrane trafficking [25], and recent studies point to a role for LRRK2 in axonal transport in polarized neurons [26-28].

Association of α-synuclein with membranes in an alpha-helical, tetramer conformation in which the NAC domain is buried protects it from forming pathologic ß-sheet aggregates [17, 29-34]. Altering the association of normal, tetrameric α-synuclein with membranes has been suggested as a therapeutic intervention against inclusion formation [30, 35-39]. G2019S-LRRK2 expression in neurons decreases membrane association of α-synuclein., suggesting a role in promoting α-synuclein aggregation [40]. Increasing neuronal activity in neurons expressing G2019S-LRRK2 reduces further exacerbates α-synuclein pathologic aggregation [41, 42]. Since α-synuclein primarily localizes to presynaptic vesicles [13, 43], it is possible that LRRK2 kinase activity plays a role in α-synuclein trafficking and localization to this compartment.

Here, we demonstrate using selective LRRK2 kinase inhibitors along with ExM, which provides exquisite resolution of synapses [44, 45], an increase in colocalization of □-synuclein with presynaptic markers. Reduction of LRRK2 kinase activity increases presynaptic targeting of α- synuclein in primary hippocampal neurons, and in glutamatergic and dopaminergic terminals in the mouse striatum. Inhibition of LRRK2 kinase activity also increases the motility and fast anterograde transport of □-synuclein along the axon. Together, our data suggest an interaction between LRRK2 kinase activity, and □-synuclein localization and targeting to presynaptic terminals. Moreover, they prompt further investigation into the mechanisms by which LRRK2 kinase activity directs vesicular transport of □-synuclein along the axon and how that may influence pathology in LRRK2-PD, and idiopathic PD.

## Materials and methods

Unless otherwise stated, all materials were purchased from Fisher Scientific.

### Animals

All animal protocols were approved by the Institutional Animal Care and Use Committee at the University of Alabama at Birmingham. Pregnant CD1 mice were purchased from Charles River (Wilmington, MA). C57BL/6J mice were from purchased from the Jackson Laboratory. PF-360 chow [46] was formulated at Research Diets Inc. Mice were on a 12-hour light/dark cycle and had ad libitum access to food and water.

### Primary hippocampal culture

Primary neuronal cultures from E16-E18 mouse embryos were generated as previously described [47]. Stocks of MLi-2 were prepared at 30 mM by dissolving in DMSO, and stored at -80°C for no more than 3 months (aliquots were not re-used). On DIV 7, neurons for immunoblotting, immunofluorescence, and ExM experiments were treated with LRRK2 inhibitor MLi-2 (10nM or 30nM respectively), or equivalent dilution of DMSO as control. Cells were lysed for western blot or fixed for immunofluorescence on DIV14.

### Live cell imaging and kymograph analyses

Primary hippocampal neurons were transfected with 1.5 μg human α-synuclein-GFP plasmid DNA [48] using lipofectamine-LTX (Thermofisher) on DIV6 and imaged on DIV10 using an inverted ZEISS Z1 Cell Observer fitted with a Colibri2 cool LED 5-channel fast wavelength switching system, and a Hammamatsu Orca Flash high-speed camera. The incubation chamber temperature was set at 32°C. Prior to recording, neuronal media was exchanged to pre-warmed imaging buffer (136 mM NaCl, 2.5 mM KCl, 2 mM CaCl_2_, 1.3 mM MgCl_2_, 10 mM glucose, 10 mM HEPES, pH 7.4) containing either 30 nM MLi-2 or DMSO control. Regions of thin axons at least 100 μm away from neurite tips, or adjacent cell bodies were chosen for imaging. The direction towards the cell soma was marked during recordings to help distinguish between anterograde (towards synaptic terminal) or retrograde (towards cell soma) direction during analysis. Images were captured every 300 ms for a total duration of 2 minutes. Kymographs were generated using the multi kymograph plugin in FIJI. Puncta count was determined by manual count within 50 μm of axon length. Mobile puncta were defined by any kind of mobility throughout the recording window. To account for bidirectional travelling puncta, directionally mobile puncta were defined as those with a net vectorial movement of 10 μm in either retrograde or anterograde direction [49]. Net velocities of anterograde and retrograde trafficking were manually measured per each track by dividing the total distance travelled by total time travelled [50]. Only mobile puncta classified as either retrograde or anterograde were used to calculate overall velocity.

### Immunoblots

Primary neurons were lysed on DIV14 using 2% SDS in Tris-buffered saline (TBS; 20mM Tris, 150 mM NaCl, pH 7.4) containing phosphatase and protease inhibitors. Cells were scraped and sonicated for a total of 10 seconds (1sec on/off pulse, 30% amplitude). Lysates were centrifuged at 20,000*g*, and the supernatant was diluted into 4X Laemmli buffer with 10% dithiothreitol. Protein lysates were electrophoresed on 8% (LRRK2) or 15% (α-synuclein) SDS-PAGE gels and transferred overnight onto PVDF membranes (Millipore). Blots were either blocked with Everyblot blocking reagent (BioRad) for LRRK2 and phospho-protein blots, or 5% milk in TBS-Tween followed by primary antibody incubation in blocking buffer overnight at 4°C. The following antibodies were used: anti-LRRK2 c42-2 (Abcam, RRID: AB_2713963), anti-LRRK2 pS1292 (Abcam, RRID: AB_2732035), anti-LRRK2 pS935 (Abcam, RRID: AB_2732035), anti-□βsynuclein (Abcam, RRID: AB_869971), anti-vGLUT1(Synaptic Systems, RRID: AB_887878), anti-VAMP2 (Synaptic Systems, RRID: AB_887811), anti-dopamine transporter N-terminus rat monoclonal (RRID: AB_2190413), and anti-vinculin (Biorad, RRID: AB_2214389). Blots were then incubated with HRP-conjugated secondary antibodies (Jackson Immunoresearch) for 2 hours at room temperature and developed with ECL (Biorad 40-720-71KIT). Blots were quantified using FIJI software.

### Immunofluorescence

#### Transcardial perfusions

C57BL/6J mice were transcardially perfused with 0.9% saline, 10 units/mL heparin, and sodium nitroprusside (0.5% w/v) followed by cold 4% paraformaldehyde (PFA), 30% acrylamide in phosphate buffered saline (PBS). After 12h post fixation in 4% PFA at 4°C, brains were embedded into a 2% agarose gel (in PBS). Agarose was dissolved in microwave-heated PBS and cooled down until 37°C and then poured into a 6-well dish containing the brains followed by solidifying at 4°C. A Leica VT1000 S was used to cut 100 μm thick brain sections which were stored in PBS at 4° C. Sections were stored no longer than 2 weeks in PBS prior to IF processing.

#### Immunofluorescence Primary neurons

Neurons were fixed with a 4% PFA, 4% sucrose in PBS for 30 min at room temperature, rinsed five times in PBS, permeabilized and blocked with 0.05% saponin, 3% BSA in PBS. This buffer was used for the primary and secondary antibody incubations. The following antibodies were used: anti-□βsynuclein (Abcam, RRID: AB_869971), anti-vGLUT1 (Synaptic Systems, RRID: AB_887878), anti-Homer1 (Synaptic Systems, RRID: AB_2631222). Primary incubations were performed at 4°C overnight followed by five rinses. Samples were incubated in Alexa Fluor-conjugated secondary antibodies (Thermofisher) for 1h at room temperature, rinsed, and mounted onto glass slides (Superfrost Plus) in Prolong Gold (Thermofisher).

#### Brain sections

Sections were rinsed three times in TBS, and then incubated in an antigen retrieval solution (10 mM sodium citrate, 0.05% Tween-20, pH 6.0) for 1 h at 37°C. Sections were blocked and permeabilized for 1h at 4°C with agitation in 5% normal goat or donkey serum, 0.1% TritonX-100 in TBS. Primary antibodies were diluted 5% normal goat or donkey serum in TBS. The following antibodies were used: anti-total-synuclein (1:2000, RRID: AB_398107), anti-vGLUT1 (1:2000, RRID: AB_887878), anti-Homer1 (1:1000, RRID: AB_2631222), anti-hDAT-NT (1:5000, RRID: AB_2190413). Sections were incubated in Alexa-Fluor conjugated secondary antibodies diluted in 5% normal serum in PBS for 2h at room temperature and mounted onto glass slides (Superfrost Plus) using Prolong gold (Thermofisher).

### Expansion microscopy

#### Primary neurons

Neurons were fixed and incubated with primary and secondary antibodies as described above. After secondary antibody incubations, neurons were anchored with succinimidyl ester of 6-((Acryloyl)amino) hexanoic acid (AcX) (Thermofisher) in PBS (0.1mg/ml) for 6h at 4°C. Samples were then incubated for 30 min at 37°C in a gelling chamber containing the polymer solution (2M NaCl, 8.625% sodium acrylate (Sigma), 2.5% acrylamide (Sigma), 0.15%N,N’-methylenebisacrylamide, 0.02% ammonium persulfate (Biorad), 0.02% TEMED (Biorad) in PBS) as described in [51] and [52]. To allow equidistant expansion, gelled coverslips were transferred into digestion solution (50 mM Tris, 1 mM EDTA, 1% Triton X-100, 0.8 M guanidine HCl, pH8) containing proteinase K (800 u/mL, NE) and incubated for 6h at room temperature. After digestion, gel-embedded neurons were transferred to glass bottom imaging plates (Cellvis) and rinsed with PBS to allow for an approximately 2.4-fold expansion of the sample followed by immediate confocal imaging. The diameters of the gelled coverslips were measured pre- and post-expansion with a ruler to calculate the expansion factor. *Brain sections:* Brain sections were obtained as described above. Polymer solution (as described above) with additional 0.02% (w/w) 4-hydroxy tempo (Sigma) was added onto brain sections in gelling chamber, followed by 30 min incubation at 4°C. In situ polymerization was then facilitated by a 2 h incubation step at 37°C. Gelled brain sections were then further dissected for regions of interests and transferred into 1.5 mL microcentrifuge tubes containing 0.5 mL digestion solution (200 mM SDS, 200 mM NaCl, 50 mM Tris, pH 9). To allow for equidistant expansion, samples were incubated for 37°C and 95°C for 1 hour, respectively. After digestion, samples were rinsed at five times with PBS, followed by a 2 h incubation in blocking solution (% donkey serum, 0.1 % Triton X-100 in PBS). Primary antibodies were diluted in 5% donkey serum in PBS and gelled samples were incubated for a minimum of 24 hours at °C [52, 53]. The following antibodies were used: anti- αβsynuclein (Abcam, RRID: AB_869971), anti-vGLUT1 (Synaptic Systems, RRID: AB_887878), anti-Homer1 (Synaptic Systems RRID: AB_2631222). ExM samples were incubated for 24h in Alexa-Fluor conjugated secondary antibodies at 4 °C. The expansion factor was calculated by measuring the dorsal-ventral and medial-lateral distances in coronal mouse brain sections stained with Hoechst 33342 for pre- and post-expansion tissue.

### Confocal microscopy and image analysis

Confocal imaging was performed on a Nikon A1R Confocal microscope with 40X and 60X oil immersion objectives. Primary neuron ExM samples in imaging plates were imaged with an inverted Nikon C2+ confocal with a galvanometer-based high-speed scanning and an apochromatic long working distance 40X water immersion objective. Brain sections were imaged as z-stacks (step size 0.2 μm) on a Nikon A1R Confocal microscope. For colocalization analysis, the coloc2 plugin for FIJI was used and the thresholded Mander’s Colocalization Coefficient (MCC) calculated. For distance measurements for ExM primary neuron experiments, synapses were identified by juxtaposed pre- and postsynaptic markers. A line perpendicular to the synaptic cleft was drawn, the grey intensity profile of each marker was plotted and the relative peak fluorescence to peak distance was measured to assess the relative distance between markers of interest. For tissue ExM experiments, maximal projection images were generated using Fiji, and synapses identified as described in [44]. After synapse identification, a line perpendicular to the synaptic cleft was drawn and the fluorescent profiles of the markers of interest used to calculate the relative distance. Experimenters conducting imaging and analysis were blinded to experimental conditions.

### Synaptosome fractionation

Three months old C57BL/6J mice received PF-360 chow or control chow for 8 days. Brains were homogenized in 2 mL of sucrose buffer (320 mM sucrose, 2 mM DDT, 1 mM EDTA, 1 mM EGTA, 4 mM HEPES·KOH, pH 7.4, with protease and phosphatase inhibitors) using a Teflon dounce on ice with 10 strokes. Differential centrifugation was used to fractionate brain homogenate as described (Lee et al., 2001). In brief, the total homogenate was centrifuged for 10 minutes (min) at 1,000*g*, the post-nuclear supernatant was centrifuged for 10 min at 9,200*g*, the resulting pellet was resuspended in sucrose buffer followed by centrifugation at 10,200*g*. The resulting pellet is the synaptosome fraction, P2. The supernatant was further centrifuged for 2h at 167,000*g* to fractionate the total soluble protein fraction (S3) and the microsomal pellet (P3) which was resuspended in 700 µL sucrose buffer and homogenized in a glass dounce with ten up-and-down strokes. The P2 pellet was resuspended in 5 mM HEPES-KOH (pH 7.4) supplemented with protease and phosphatase inhibitors, homogenized in with a glass dounce and centrifuged for 20 min at 21,130*g*. The resulting synaptosomal, membrane-enriched pellet (LP1) was resuspended in sucrose buffer and the supernatant was centrifuged for 2h at 200,000*g*. Finally, the sedimented synaptic vesicle-enriched fraction (LP2) was resuspended sucrose buffer and homogenized with a glass dounce. All isolated fractions were snap-frozen in EtOH/dry ice slurry and stored at □80 °C until use.

### Statistics

All statistical analyses were performed using Graph Pad software. Data were presented as mean and standard error of the mean unless indicated differently in the figure legends. Nested independent t-test analysis or nested one-way ANOVA with Tukey’s posthoc analyses were performed. If data did not fit a normal distribution, the Mann-Whitney test was used. Statistical analyses are presented in Table 1.

## Results

### LRRK2 kinase inhibition increases colocalization of α-synuclein and the presynaptic marker vGLUT1 in primary hippocampal neurons

α-Synuclein is highly expressed in glutamatergic presynaptic terminals where it colocalizes with vGLUT1 [15, 54]. Furthermore, LRRK2 is also expressed in excitatory corticostriatal neurons [55], where its kinase activity influences striatal excitatory transmission [56-58]. To determine if LRRK2 kinase activity affects presynaptic localization of α-synuclein, primary hippocampal neurons, which are predominantly glutamatergic, were treated with the LRRK2 kinase inhibitor MLi-2, a selective LRRK2 kinase inhibitor [59], or DMSO vehicle control for 7 days [10]. Double labeling immunofluorescence and confocal microscopy were performed for α-synuclein and vGLUT1. In control primary hippocampal neurons, α-synuclein colocalized with vGLUT1 at excitatory presynaptic terminals (Figure 1A). In neurons treated with MLi-2 for 7 days, colocalization of α-synuclein and vGLUT1 significantly increased compared to control neurons (Figure 1A, B). Immunoblots of primary hippocampal neuron lysates treated with DMSO control or MLi-2 (10 nM, 30 nM) using an antibody that selectively recognizes the LRRK2 autophosphorylation site, S1292 [60], showed that MLi-2 significantly reduced LRRK2 kinase activity. Expression levels of α-synuclein and vGLUT1 were not significantly altered upon 7 days of LRRK2 kinase inhibition (Figure 1C, D). In addition to unaltered protein expression of the glutamatergic terminal marker vGLUT1 upon MLi-2 treatment, inhibition of LRRK2 kinase activity did not alter the overall density of glutamatergic terminals in primary hippocampal cultures (Figure S1). These findings suggest that LRRK2 kinase activity increases the localization of α-synuclein at the presynaptic terminal.

**Figure 1:**
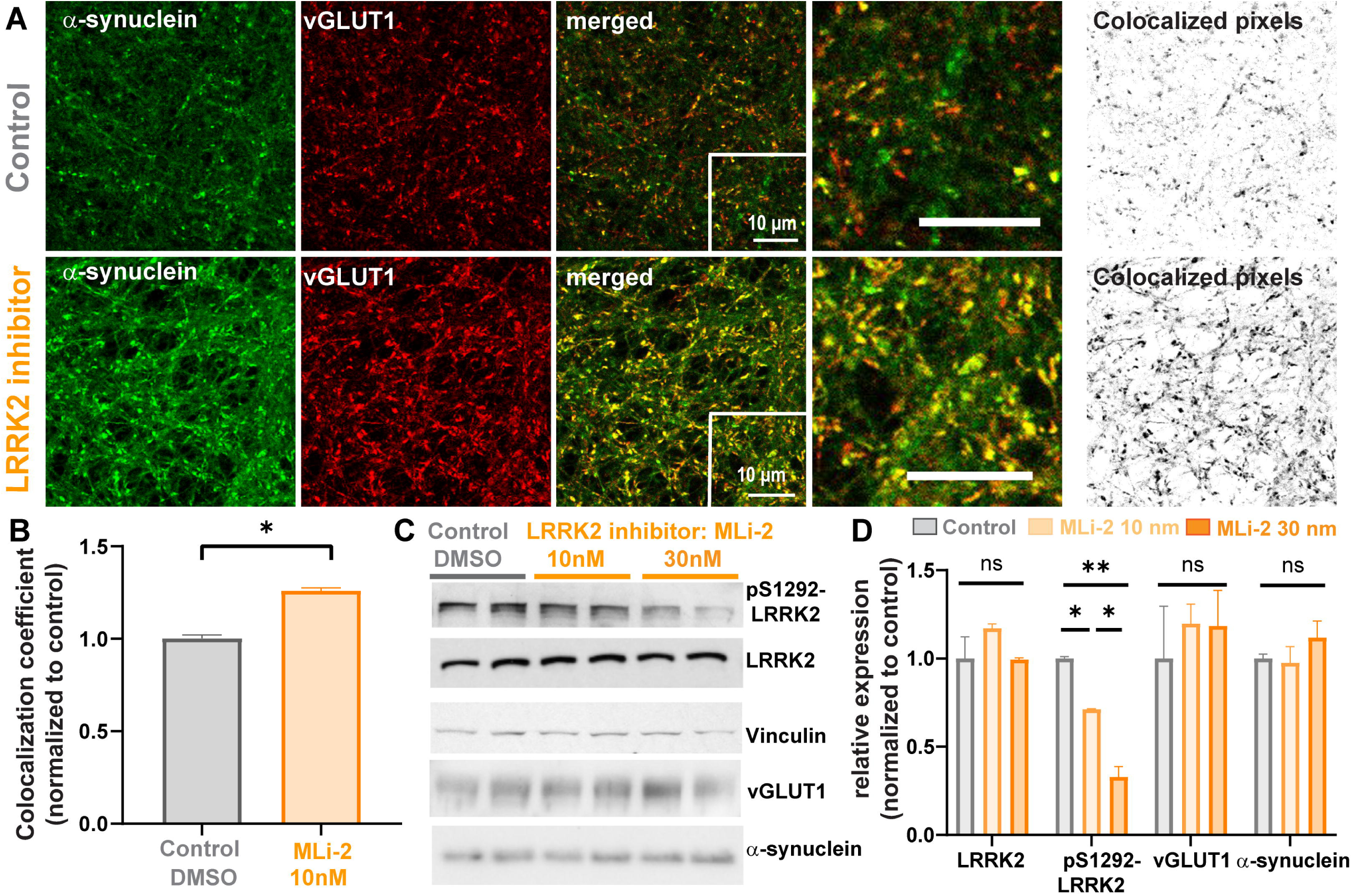
LRRK2 inhibition increases colocalization of α-synuclein with glutamatergic presynaptic marker, vGLUT1, in primary hippocampal neurons. **A)** Confocal microscope images of α-synuclein (green) and vGLUT1 (red) in primary hippocampal neurons. The upper panel shows primary hippocampal neurons after 7 days control treatment (DMSO at equivalent dilution as MLi-2); the lower panel shows primary hippocampal neurons treated for 7 days with 10 nM MLi-2. The images on the right side represent all colocalized pixels (black) between α-synuclein and vGLUT1 immunostainings generated using Fiji software. Scale bar = 10 µm. **B)** Quantification of colocalization analysis. MCCs between α-synuclein and vGLUT1 were normalized to the control treatment. Nested t-test analysis showed significant difference between treatments (N=3, 30 measurements per sample). t(6) = 2.9. *p<0.05. **C)** Immunoblots from two independent samples of primary hippocampal neurons treated with DMSO control, 10 nM MLi-2 or 30 nM MLi-2 for 7 days. **D)** Quantification of immunoblots (n=2). Shown are the normalized mean values (vinculin as loading control) of the relative expression of LRRK2, pS1292-LRRK2, vGLUT1 and α-synuclein. One-way ANOVA revealed significant inhibition of LRRK2 autophosphorylation (pS1292-LRRK2) for 10nM and 30nM MLi-2 treatment. F(2,3) = 9.5. *p<0.05; **p<0.01.

### LRRK2 kinase inhibition increases α-synuclein overlap with presynaptic vGLUT1

LRRK2 localizes both pre- and post-synaptically in the rodent brain [55]. ExM was used to resolve the presynaptic terminal, postsynaptic densities, and localization of α-synuclein. ExM overcomes resolution limitations of traditional light microscopy by physically expanding the sample [51, 52, 61]. In traditional confocal images, α-synuclein colocalized with presynaptic vGLUT1, but both also showed overlap with the post-synaptic density marker, Homer1 (Figure 2A). By using ExM, the resolution of confocal images was increased by 2.4 X, allowing presynaptic vGLUT1 and α-synuclein to be visibly distinguished from postsynaptic Homer1 (Figure 2B). In primary hippocampal neurons treated with 10 nM MLi-2 for 7 days, α-synuclein showed increased colocalization with vGLUT1 (Figure 1C, D). The distance between the maximal fluorescence intensity peaks [44, 61] of Homer1 and vGLUT1 in control neurons was approximately 0.28 µm and was not significantly altered in MLi-2 treated neurons compared to control treated neurons, indicating no change in the distance across the synaptic cleft (Figure 2B-D). The distance between the maximal fluorescence intensity peaks of α-synuclein and vGLUT1 was significantly reduced in MLi-2 treated neurons (average of 0.04 µm) compared to control neurons (average of 0.12 µm), indicating increased overlap of α-synuclein and presynaptic vGLUT1 (Figure 2B-D). These data demonstrate that increased overlap of α-synuclein with vGLUT1 at the presynaptic terminal is not a result of altered pre- and postsynaptic morphology in general, rather a redistribution of α-synuclein caused by reduced LRRK2 kinase activity. In addition, these experiments confirm that unlike traditional confocal microscopy, ExM provides a method of quantifying changes in protein localizations in small, subcellular compartments.

**Figure 2:**
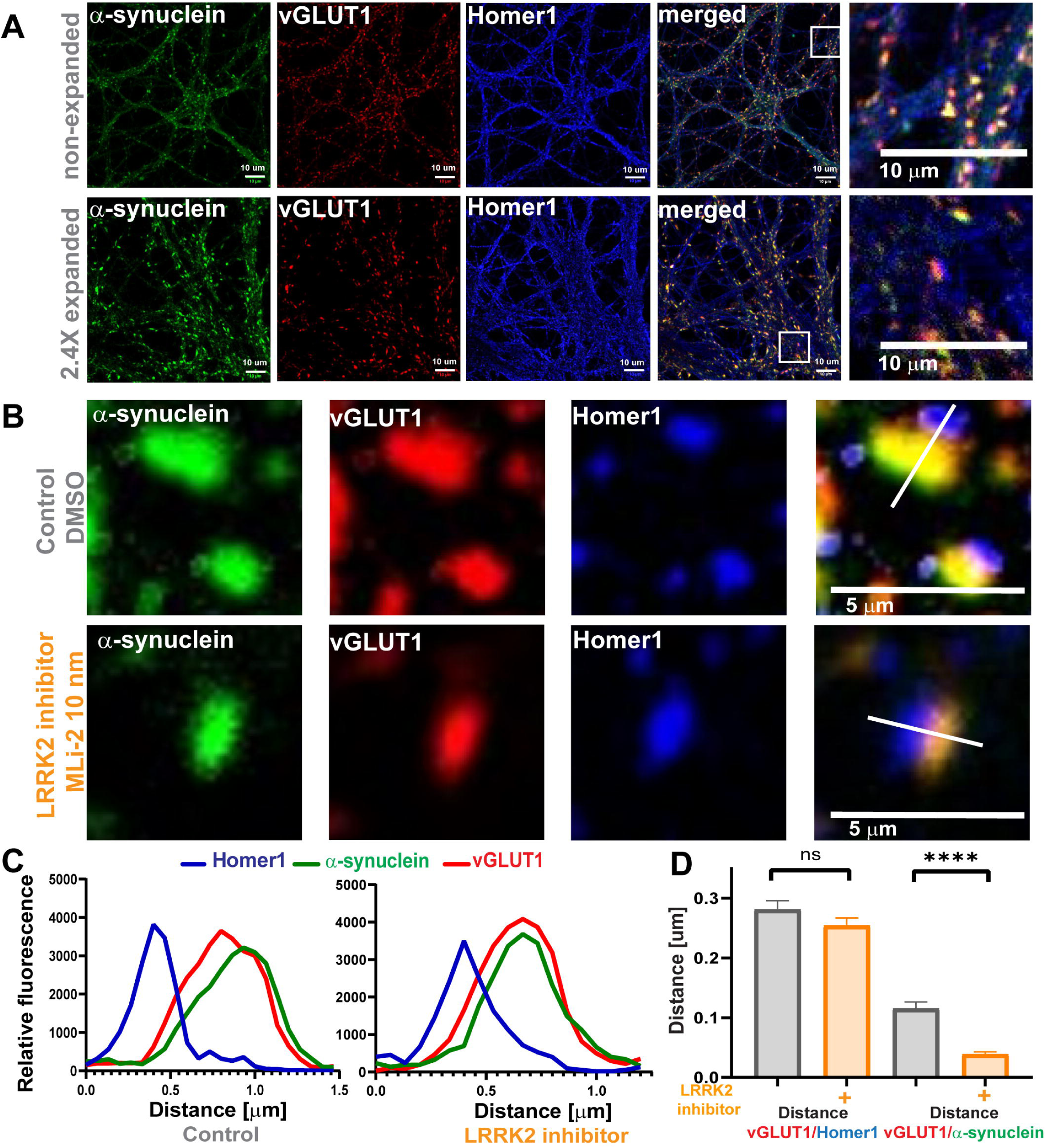
LRRK2 inhibition reduces relative distance between α-synuclein and vGLUT1 in primary hippocampal neurons. **A)** Confocal microscope images of non-expanded and 2.4 expanded primary neurons immunostained for α-synuclein (green), vGLUT1 (red) and Homer1 (blue). Scale bar= 10 µm. **B)** Representative synapses from 7 days treatment with DMSO control and LRRK2 kinase inhibitor treatment (10 nM MLI-2 Scale bar = 5 µm.) **C) D)** Quantification of distance measurements. (Nested t-test t = (116) = 6.5. ****p<0.0001, n=2, 25 measurements per sample).

### LRRK2 kinase inhibition increased α-synuclein motility and anterograde trafficking

A portion of α-synuclein associates with membranes traveling via fast axonal transport in both anterograde and retrograde directions [48]. To determine if increased anterograde axonal transport at least partially accounts for enhanced localization of α-synuclein at the presynaptic terminal in response to LRRK2 kinase inhibition, we transfected cells with α-synuclein-GFP and imaged axonal transport using live cell microscopy. Neurons were treated with 30 nM MLi-2 30 minutes prior to imaging. Figure 3A shows examples of axons containing α-synuclein-GFP puncta and representative kymographs demonstrating puncta movement over time. Pharmacological reduction of LRRK2 kinase activity with MLi-2 treatment significantly increased the percentage of mobile α-synuclein carriers moving in the anterograde direction, but did not impact the percentage of mobile carriers in the retrograde direction (Figure 3B). Non-directional mobile puncta were classified by moving puncta that did not reach the net vectorial distance of 10 μm in either direction. The percentage of mobile α-synuclein-GFP was significantly increased (Figure 3B). The overall abundance of α-synuclein puncta was unaltered. The overall net velocities of anterogradely traveling α-synuclein-GFP puncta were not significantly different between control and MLi-2 treated neurons (Figure 3D). Analyses of binned velocities of α-synuclein-GFP particles (see [40]) revealed a shift toward faster velocities relative to the control neurons (Figure 3C), however the differences in velocities between control and MLi-2 treated neurons were not significant. Overall, these data indicate that LRRK2 kinase inhibition increases movement of membranes bearing α-synuclein toward the presynaptic terminal.

**Figure 3:**
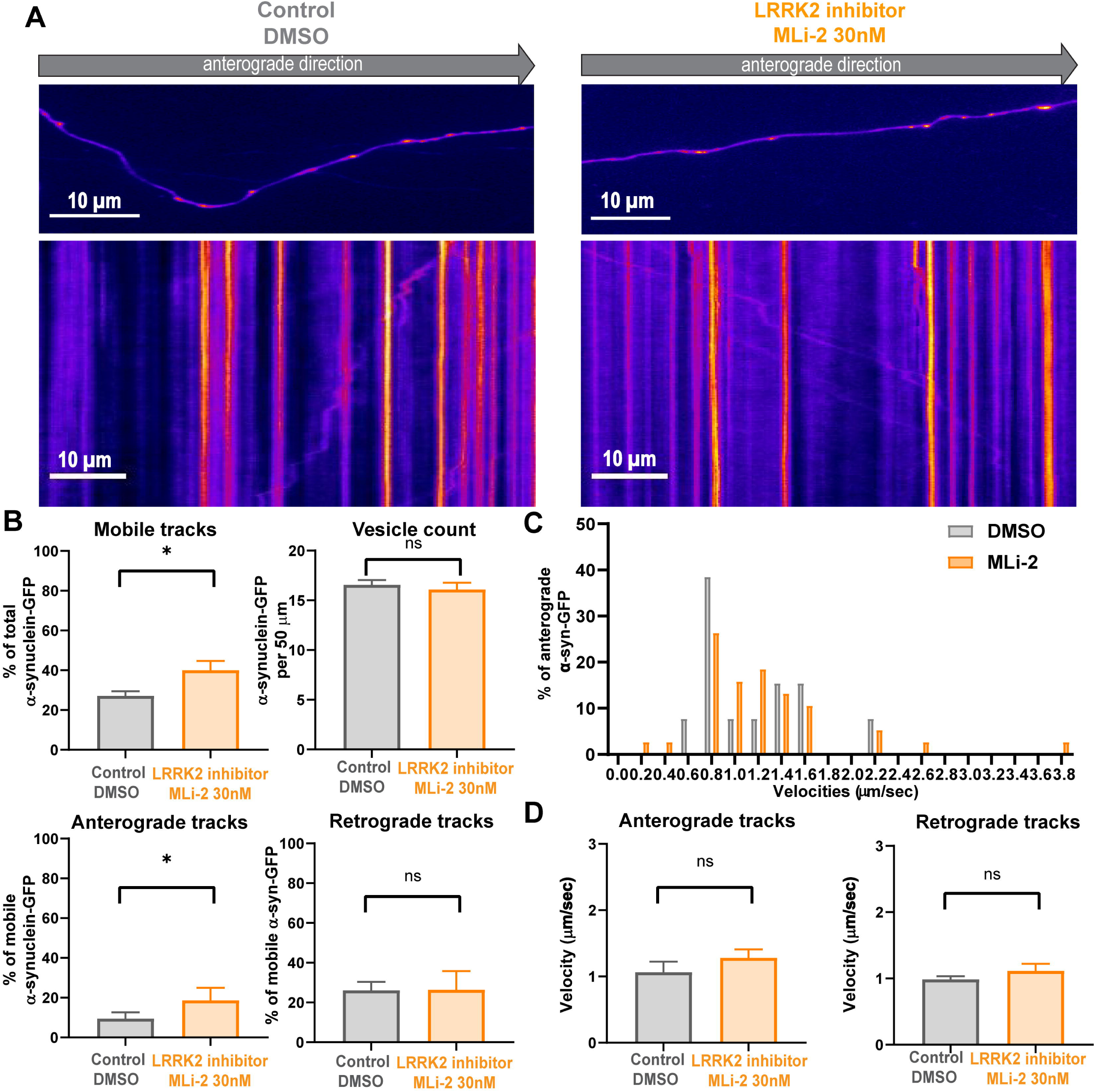
LRRK2 inhibition increases α-synuclein mobility and percentage of α-synuclein-GFP undergoing anterograde axonal transport. **A)** Primary hippocampal neurons were transfected with human α-synuclein-GFP on DIV10, treated with 30 nM MLi-2 or equivalent dilution of DMSO 30 minutes before live cell imaging. Shown are representative snapshots of the axons expressing α-synuclein-GFP and their respective kymographs showing transport over time. Images were captured every 300 ms for 2 minutes. Scale bar=10 µm. **B)** Quantification of mobile tracks (traveled ≥ 10 µm) (Nest t test, t(6)= 6.5, *p<0.05, α-synuclein-GFP puncta count per 50 µm axon length (t(6) = 0.6), and percentages of anterograde (t(51) = 2.2 ; *p<0.05) and retrograde tracks of mobile puncta(t(6) = 0.5) (N=4, number of analyzed kymographs: 27 (DMSO control), 26 (30nM MLi-2)). **C)** Distribution of binned velocities of anterograde traveling α-synuclein-GFP in neurons treated with DMSO or MLi-2. **D)** Quantification of mean velocities for anterograde (t (6) =0.4) and retrograde α-synuclein-GFP (t (87) =1.8). (N=4, number of analyzed kymographs: 27 (DMSO control), 26 (30nM MLi-2)).

### LRRK2 and α-synuclein are enriched at presynaptic vesicle membranes in vivo

To determine if LRRK2 kinase inhibition affects α-synuclein localization *in vivo*, an in-diet dosing approach for chronic LRRK2 inhibition was used as described previously [46]. C57BL/6J were fed chow formulated with 175 or 350 mg kg^-1^ PF-360 for 7 days to achieve chronic LRRK2 kinase inhibition. Reductions in LRRK2 kinase activity were assessed by immunoblotting for pS935-LRRK2 and total LRRK2 protein to determine the ratio of phospho-to-total LRRK2 in forebrain homogenates. pS935-LRRK2 levels are considered an indirect measure of LRRK2 kinase activity [46, 62, 63]. Both concentrations of PF-360 chow demonstrated reduced levels of pS935-LRRK2 without reducing total LRRK2 protein levels (Figure 4A), as previously shown [46].

**Figure 4:**
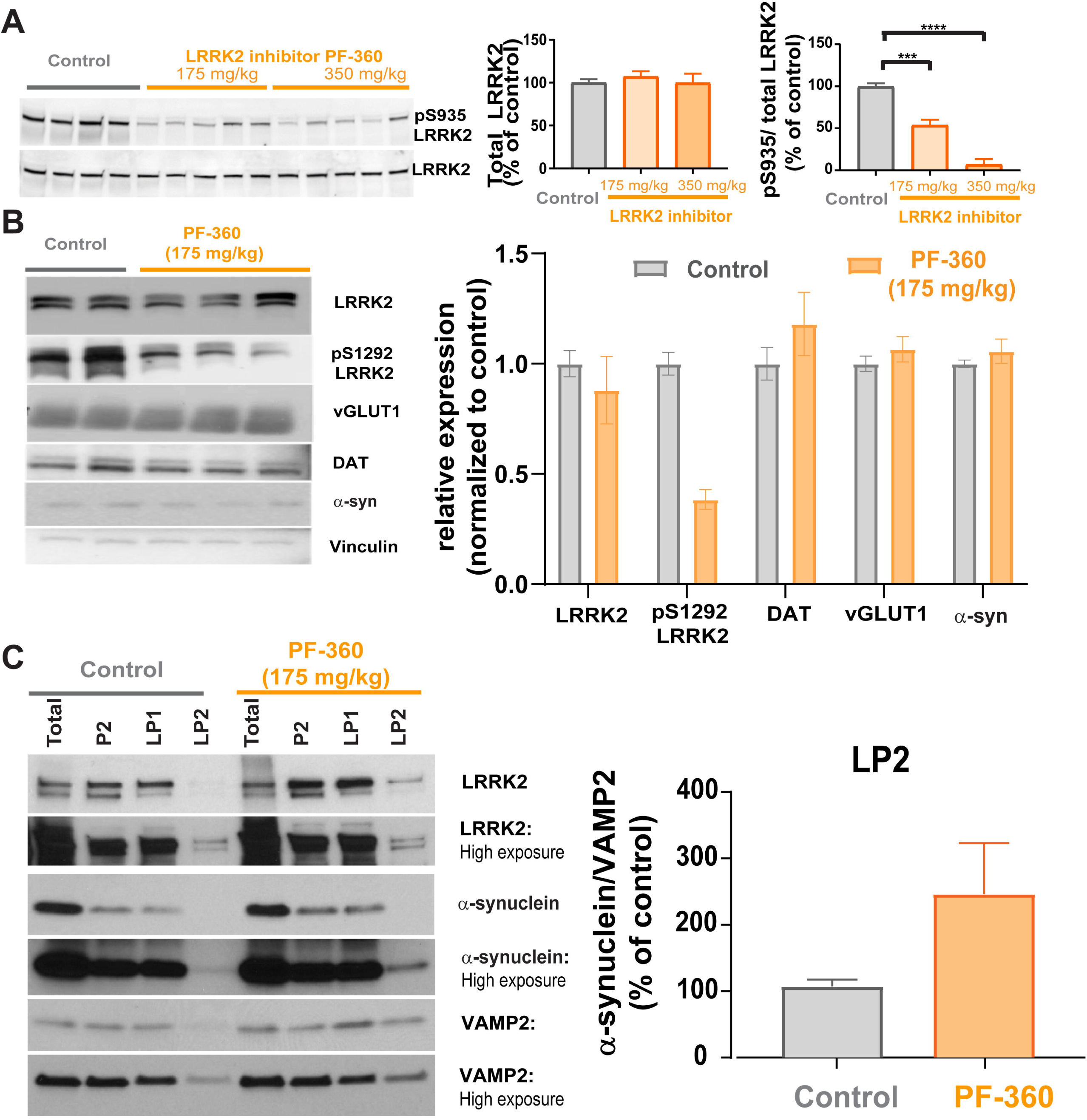
Inhibition of LRRK2 kinase activity in vivo and synaptosomal presynaptic distribution of LRRK2 and α-synuclein. **A)** C57BL/6J mice (3 months of age) received 7 days access to control chow (n=4) or chow with 175mg/kg (n=5) or 350 mg/kg (n=5) PF-360. Forebrain homogenates were immunoblotted using antibodies to pS935-LRRK2 or total LRRK2. Total LRRK2 was quantified by normalizing to total protein on the stain free Biorad gel imaged on Chemdoc. Data is expressed as an average percentage (+/-SEM) of LRRK2 from control mice. pS935-LRRK2 was quantified by normalizing the signal to total LRRK2 and data is expressed as an average percentage (+/- SEM) of LRRK2 from control mice. One-way ANOVA revealed significant inhibition of LRRK2 kinase activity. F (2, 13)=5.6. ***p<0.001, ****p<0.0001. **B)** A separate cohort of C57BL/6J mice were given access to control chow (n=2) or 175 mg/kg PF-360 chow (n=3) for 7 days. Whole brain lysates were immunoblotted for LRRK2, pS1292-LRRK2, presynaptic vGLUT1 and DAT, α-synuclein, and loading control vinculin. Relative protein expression was quantified by normalizing to (vinculin) followed by normalization to control treatment and is indicated as mean values (+/- SEM). pS1292-LRRK2 levels were reduced upon PF-360 treatment. **C)** Forebrains were fractionated into total homogenate, P2 (crude synaptosomes), LP1 (synaptosome membrane fraction) and LP2 synaptic vesicle enriched fraction. Fractions were immunoblotted for LRRK2, α-synuclein, VAMP2. The right panel shows quantitation of α-synuclein relative to presynaptic VAMP2.

A separate cohort of mice fed either control or PF-360 chow (175mg/kg) for 7 days was processed for whole brain lysates for immunoblot analyses of synaptic markers of interest for this study. In addition to synaptic marker and α-synuclein immunoblotting, the levels of the direct LRRK2 kinase phosphorylation site pS1292-LRRK2 were also assessed (Figure 4B). While PF-360 treatment reduced levels of pS1292-LRRK2, quantification did not reveal a statistically significant change. Levels of presynaptic markers vGLUT1 and dopamine transporter (DAT) were not changed upon PF-360 treatment, nor did α-synuclein levels change upon LRRK2 kinase inhibitor treatment.

LRRK2 localizes to and plays a functional role at the presynaptic terminal [18, 20, 21, 23, 64, 65]. In addition, α-synuclein has been shown to be enriched at glutamatergic terminals [15, 54]. Synaptosome fractionation of forebrain homogenates in a separate cohort of C57BL/6J mice were used to determine whether 7 days of treatment with PF-360 altered localization of α-synuclein in the synaptic vesicle fraction. First, we found that LRRK2 localized to the synaptic vesicle enriched fraction LP2, confirming presynaptic LRRK2 (Figure 4C). Immunofluorescence co-labelling for LRRK2 and presynaptic marker vGLUT1 and VAMP2 revealed colocalized puncta in the mouse brain striatum (Figure S2), further demonstrating LRRK2 localization glutamatergic presynaptic terminals. LRRK2 KO striatum showed minimal LRRK2 immunofluorescence signal (Figure S2), demonstrating specificity of our LRRK2 immunofluorescence results.

Seven days of treatment with PF-360 increased the proportion of α-synuclein in the LP2 fraction relative to controls, consistent with our imaging experiments in primary hippocampal neurons. These data show that LRRK2 and α-synuclein localize to the presynaptic terminal in the striatum *in vivo*.

### LRRK2 kinase inhibition increases overlap of α-synuclein and presynaptic markers in the mouse striatum *in vivo*

To assess whether LRRK2 kinase inhibition increases presynaptic localization of α-synuclein in cortico-striatal and nigral-striatal presynaptic terminals, immunofluorescence and confocal microscopy for α-synuclein and vGLUT1, or DAT were performed in striatal sections from mice fed PF-360 (175mg/kg) or control chow for 7 days (Figure 6A). In control mice, the majority of α-synuclein overlapped with vGLUT1 as demonstrated previously [15]. Upon LRRK2 kinase inhibition, the overlap of α-synuclein with vGLUT1 significantly increased, similar to the findings in primary excitatory neurons. A smaller portion of □-synuclein colocalized with DAT (Figure 5B). LRRK2 kinase inhibitor treatment increased colocalization of α-synuclein and DAT. Thus, LRRK2 kinase inhibition also increases α-synuclein presynaptic localization in the striatum *in vivo*. Immunoblotting analysis for whole brain lysates of rodents fed PF-360 showed that LRRK2 kinase inhibition did not change the overall protein expression levels of α-synuclein, vGLUT1, or DAT (Figure 4B).In addition, LRRK2 kinase inhibition did not alter the density of glutamatergic terminals in the mouse brain striatum of animals fed PF-360 compared to control chow animals (Figure S1). The integrated density of DAT signal in the dorsal mouse striatum was significantly increased upon LRRK2 kinase inhibition comparted to control animals (Figure S1). To increase the resolution of α-synuclein, vGLUT1, and Homer1 in the striatum, we performed ExM of striatal mouse brain sections and analyzed individual synapses for the subcellular localization of α-synuclein relative to the presynaptic marker. ExM allowed resolution of synaptic markers in the mouse brain striatum compared to non-expanded samples (Figure 6A). The results of individual synapse analyses utilizing ExM revealed a significantly reduced overall distance between α-synuclein and vGLUT1 at the presynaptic terminal of glutamatergic neurons projecting into the mouse striatum for animals treated with PF-360 (Figure 6C, D). In addition, no change in the distance between Homer1 and vGLUT1 was observed upon LRRK2 kinase inhibition, and thus morphological synaptic structural rearrangements do not account for the reduced distance between α-synuclein and vGLUT1. No effect of LRRK2 kinase inhibition on the localization of the presynaptic protein synapsin-1 at glutamatergic terminals in the striatum was observed (Figure S3). Overall, our data demonstrate in primary neurons in culture and in mouse brains in vivo that reduced LRRK2 kinase activity increases presynaptic targeting of α-synuclein.

**Figure 5:**
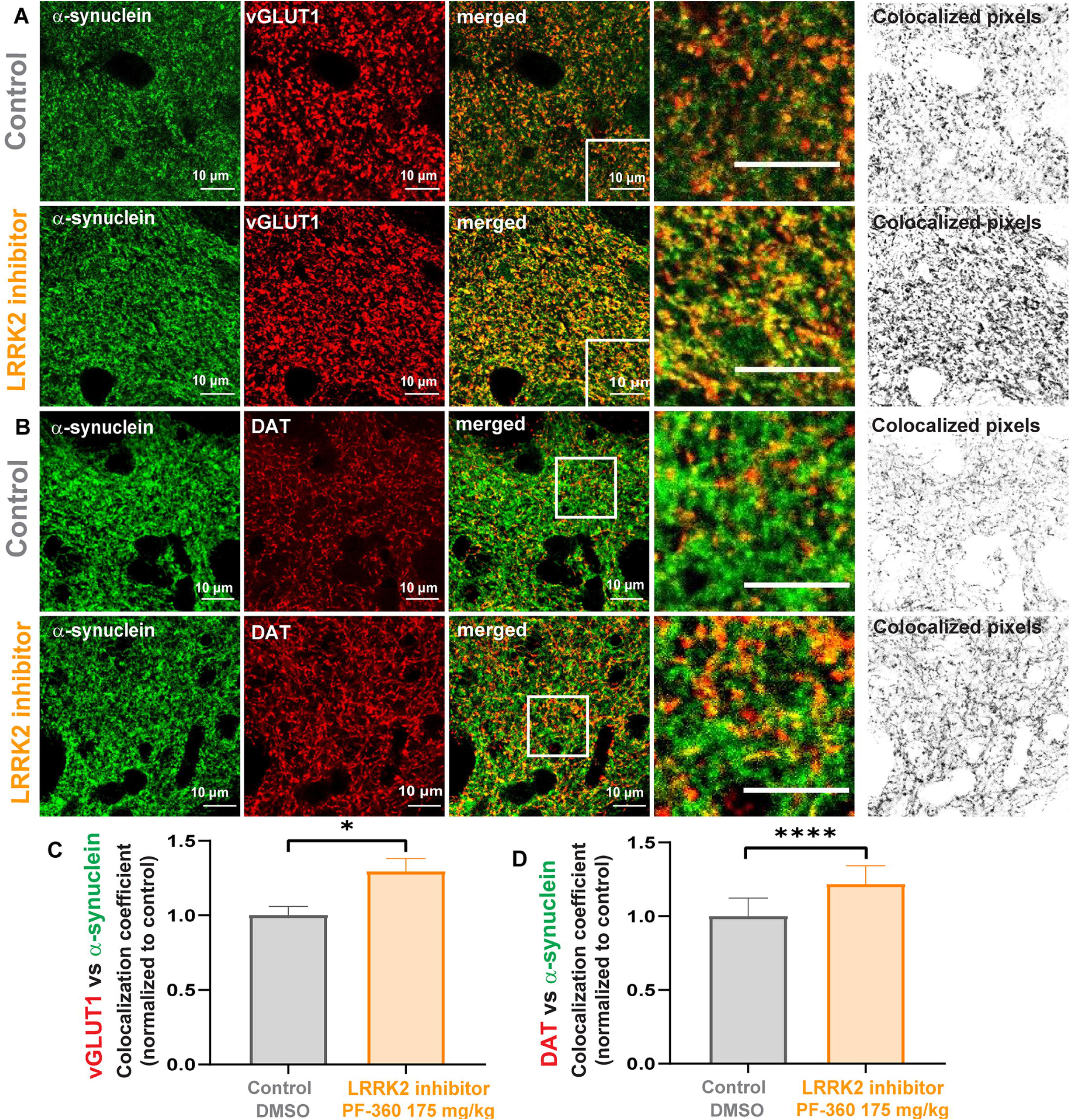
LRRK2 kinase inhibition increases colocalization of α-synuclein and presynaptic markers in the striatum. **A)** Confocal microscope images of α-synuclein (green) and vGLUT1 (red) in the dorsal striatum in mice fed with control (top panels) or 175 mg/kg PF-360 (lower panels) for 7 days. The images on the right represent all colocalized pixels (black) between α-synuclein and vGLUT1 immunostainings. Scale bar = 10 µm. **B)** Confocal microscope images of α-synuclein (green) and dopaminergic terminal marker DAT (red) in the mouse dorsal striatum. The top panels are from control mice and the lower panels are from the mice fed PF-360 chow for 7 days. The images on the right represent all colocalized pixels (black) between α-synuclein and DAT immunostainings. Scale bar = 10 µm **C)** Quantification of colocalization analysis. MCCs between α-synuclein and vGLUT1, and α-synuclein and DAT were normalized to control treatments and are indicated as mean values (+/- SEM). Nested t-test (t (6) = 3.3 showed significance (n=3, for α-synuclein/vGLUT1, n=3 (t (178) = 6.6) for α-synuclein/DAT, with 30 individual measurements per n). *p<0.05, ****p<0.0001.

**Figure 6:**
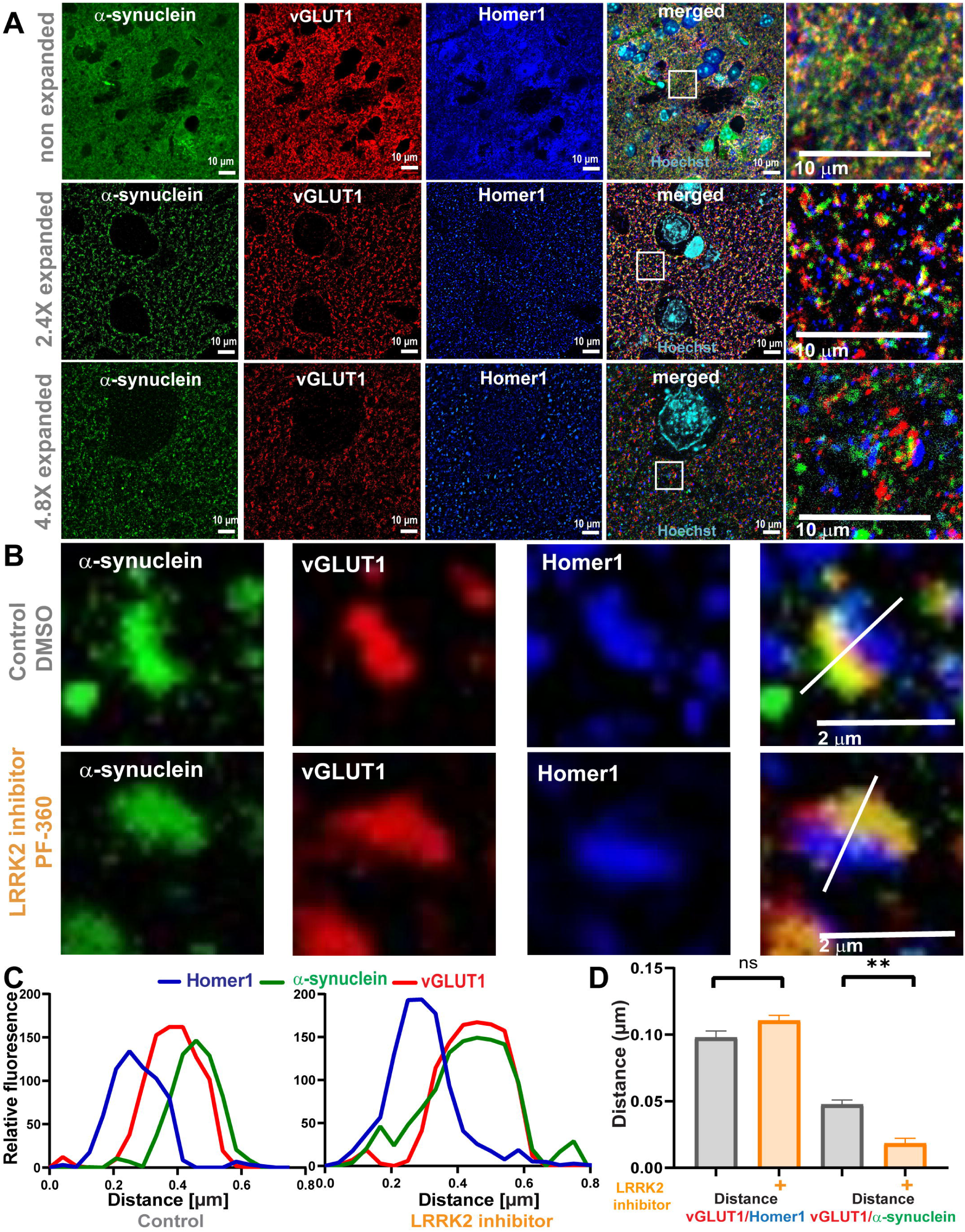
LRRK2 inhibition reduces relative distance between α-synuclein and vGLUT1 in the mouse striatum. **A)** Confocal microscope images of the striatum in non-expanded, 2.4x, and 4.8x expanded brain sections immunostained for α-synuclein (green), vGLUT1 (red) and Homer1 (blue). Scale bar = 10 µm. **B)** Representative synapses upon control and LRRK2 kinase inhibitor treatment (7 days, 10 nM MLi-2). Scale bar = 2 µm. **C)** Respective fluorescent profiles from B. **D)** Quantification of distance measurements in mean distance (+/- SEM) between Homer1 and vGLUT1 (t(4) = 0.6) and between vGLUT1 and α-synuclein (t(4) = 5.3 (nested t-test analysis, n=3, 25 measurements per n. ** p<0.01.

## Discussion

The data in this study reveal that inhibition of LRRK2 kinase activity alters the axonal transport and presynaptic localization of α-synuclein in neurons. ExM provided a method to expand the tissue to more precisely resolve and quantify proximity between α-synuclein and synaptic markers. Inhibition of LRRK2 kinase activity in both primary neurons *in vitro* and in mouse striatum *in vivo* significantly increased colocalization of α-synuclein with presynaptic markers. Reduced LRRK2 kinase activity increased localization of α-synuclein with presynaptic vGLUT1 in excitatory terminals, and overlap of α-synuclein and DAT in dopamine axons. Lastly, LRRK2 kinase inhibition also increased the motility and anterograde transport of α-synuclein in axons. We previously demonstrated that neuronal expression of G2019S-LRRK2 with elevated kinase activity increases the cytosolic fraction of α-synuclein, and multiple studies show G2019S-LRRK2 increases the abundance of pathologic α-synuclein inclusions [6-11, 17, 66]. Thus, together, these data suggest that LRRK2 kinase activity may influence the ability of α-synuclein to form pathologic aggregates by controlling the presynaptic targeting of α-synuclein.

There are several mechanisms by which LRRK2 may convert normal α-synuclein to a conformation that forms pathologic fibrils, including impairment of chaperone-mediated autophagy, or increased synthesis [67, 68]. LRRK2 also plays a major role in membrane trafficking [25, 69]. Defects in trafficking and targeting of α-synuclein to the correct membrane compartment within the neuron may influence its propensity to aggregate [70]. α-Synuclein primarily localizes to the presynaptic terminal [16, 71-74] where it interacts with the SNARE complex to reduce synaptic vesicle exocytosis [13, 14]. α -Synuclein preferentially associates with synaptic vesicle membranes and lipids as multimers, which protect the protein from forming ß-sheets, and eventually inclusions [29-32]. We previously showed using fluorescence recovery after photobleaching that expression of G2019S-LRRK2 increases the cytosolic fraction of α-synuclein [10]. The current study demonstrated that inhibition of LRRK2 kinase activity increases the normal localization of α-synuclein at the presynaptic terminal. Therefore, in addition to LRRK2-mediated alterations in proteostasis, the effect of LRRK2 on α-synuclein subcellular localization may be another mechanism by which it influences formation of pathologic inclusions.

LRRK2 and α-synuclein are highly expressed in excitatory neurons, [55]. LRRK2 kinase activity influences the physiological activity of excitatory neurons [55, 58, 75, 76]. High expression levels and the presynaptic enrichment of α-synuclein at glutamatergic terminals may have crucial implications for glutamate transmission [77, 78]. The interaction of α-synuclein and VAMP2 at excitatory terminals is thought to act as a break on vesicle release [79]. Because mutant LRRK2 increases corticostriatal excitatory transmission [57, 58] increased targeting of a-synuclein to the presynaptic terminal by LRRK2 kinase inhibitors may help dampen and rescue this aberrant activity [47]. Also, dopaminergic neurons are of major importance due to their selective vulnerability in PD. We showed that inhibition of LRRK2 kinase increases the overlap of α-synuclein and DAT in vivo, and observed an increase of DAT signal in the striatum of LRRK2 kinase inhibitor treated animals. Future experiments should elucidate the functional interaction of LRRK2 and α-synuclein in dopaminergic neurons.

To further elaborate, our data shows that inhibition of LRRK2 kinase activity increases localization of α-synuclein at the presynaptic terminal *in vitro* and *in vivo*. Our ExM data showed reduced proximity between □-synuclein and vGLUT1 at the presynaptic terminal upon LRRK2 kinase inhibition. Although the functional implications of increased targeting of α-synuclein to the presynaptic terminal are unknown, because α-synuclein can act as a brake on synaptic vesicle exocytosis it is possible that increased synaptic vesicle association of α-synuclein could reduce excitatory transmission [74, 78, 80, 81]. Previous studies using electrophysiological recordings in the striatum showed that increased LRRK2 kinase activity increases spontaneous evoked postsynaptic current frequency, suggesting increased cortico-striatal presynaptic activity. The increased cortico-striatal presynaptic activity is reversed by LRRK2 inhibition [20, 58, 64]. Cortico-striatal glutamatergic transmission and activity has been associated as a putative somatotopic stressor of dopaminergic neurons [82]. Thus, it is possible that mutant LRRK2 through its presynaptic actions on α-synuclein localization and glutamatergic synapse activity could cause a degeneration in dopamine terminals in the striatum, ultimately leading to Parkinson’s disease. Although LRRK2 inhibition did not affect localization of synapsin-1, future studies should also investigate if the observed impact of LRRK2 kinase inhibition is specific for α-synuclein, or if other synaptic proteins might be affected [83]. Electron microscopy studies show that reduced levels ofLRRK2 decreased the number of docked vesicles at the active zone, suggesting reduced LRRK2 activity could impact synaptic proteins involved in docking [84].

We also found that LRRK2 kinase inhibition increases α-synuclein colocalization with DAT in the dorsal striatum. A synergistic interaction of DAT and α-synuclein has been previously reported, as DAT activity is thought to increase membrane association of α-synuclein which in turn targets DAT to cholesterol-rich membrane domains [85]. Pathological LRRK2 mutations have been shown to reduce dopamine transporters in the striatum for patients with and without a PD diagnosis [86]. Our data showed increased DAT signal in the dorsal striatum of animals treated with LRRK2 kinase inhibitor, while overall expression of DAT was unaltered, indicating re-localization or trafficking of DAT. DAT has also been shown to mislocalize in PD post mortem brain tissue, where close proximity of DAT to aggregate-like α-synuclein has been reported [87]. Thus, the data in this study point toward a possible critical link between LRRK2 kinase activity and α-synuclein in regulating DAT activity and trafficking. Further experiments will be needed to investigate the spatial relationships and functional interactions of LRRK2, α-synuclein, and DAT.

Our data suggest that LRRK2 kinase inhibition increases α-synuclein axonal transport toward the presynaptic terminal. Although slow component B, a transport mechanism mediating axonal trafficking of soluble proteins, is thought to be the primary driver of cytosolic α-synuclein transport [88], fast axonal transport of α-synuclein on vesicles also occurs [48]. In our control neurons, there were few α-synuclein particles traveling via fast axonal transport, but this significantly increased upon LRRK2 kinase inhibition. Additionally, overall motility of α-synuclein-GFP puncta was significantly increased upon LRRK2 kinase inhibitor treatment. LRRK2 kinase inhibition significantly decreased the percentage of immobile puncta. Our data contribute to multiple emerging studies demonstrating a functional role for LRRK2 in microtubule-based axonal transport [26-28, 89]. Expression of the R1441C-LRRK2 or Y1699C-LRRK2 mutants inhibit axonal transport of mitochondria, which can be rescued by increasing microtubule acetylation [28]. The active conformation of LRRK2 associates with microtubules and blocks movement of kinesin and dynein motors [90]. Impaired balance of the kinesin and dynein motors by overactive LRRK2 reduces the processivity of autophagosome transport along the axon. Rab3a, an LRRK2 kinase substrate, mediates the interaction with kinesin-3 [91]; the active conformation of Rab3a promotes its interaction and consequently, overall anterograde trafficking of synaptic vesicle precursors in the axon. Importantly, LRRK2 kinase mediated phosphorylation of Rab GTPases is thought to disrupt their interactions with effectors [92-95]. In this way, it is conceivable that increased LRRK2 kinase activity might disrupt the transport mechanisms involving vesicles, Rabs, and associated motor proteins. Conversely, by inhibiting LRRK2 kinase activity, vesicle interactions and anterograde trafficking are promoted.

This study sheds light on the different roles LRRK2 might play in α-synuclein trafficking. These findings are consistent with previous studies demonstrating a major role of LRRK2 in membrane trafficking, and in axonal transport. Understanding LRRK2 kinase activity could be potentially important for putative treatment options of both LRRK2-PD and sporadic PD cases [4], Given that tetrameric, native α-synuclein is resistant to aggregation [30, 38, 39], our data also suggest that LRRK2 kinase inhibitors could have a therapeutic benefit in halting Parkinson’s disease progression and development of nonmotor symptoms by increasing native, presynaptic α-synuclein at membranes and protecting against pathologic aggregation.

## Conclusions

Both LRRK2 and α-synuclein localize to the presynaptic terminal, and LRRK2 kinase activity has been shown by multiple studies to impact pathologic α-synuclein aggregation, but how LRRK2 kinase influences normal α-synuclein has remained unclear. This study shows that inhibition of LRRK2 kinase activity increases presynaptic targeting of α-synuclein. Treatment with the selective LRRK2 inhibitors MLi-2 and PF-360 in primary neurons *in vitro* and in the mouse striatum *in vivo* increased overlap of α-synuclein and glutamatergic terminal marker, vGLUT1. In addition, using the high resolution imaging technique expansion microscopy to resolve and quantify changes in α-synuclein presynaptic localization, we show that inhibition of LRRK2 kinase increases the overlap of α-synuclein and vGLUT1. Live cell imaging experiments showed increased motility and anterograde trafficking of α-synuclein-GFP puncta upon LRRK2 kinase activity, consistent with growing evidence for a role of LRRK2 in axonal transport. Overall, this study demonstrate a role of LRRK2 kinase activity in α-synuclein localization at the presynaptic terminal which suggest a role for these two proteins in synaptic dysfunction and PD-related phenotypes.

## List of Abbreviations

DAT: Dopamine transporter
ExM: Expansion microscopy
LRRK2: Leucine-rich repeat kinase 2
MCC: Mander’s Colocalization coefficient
PD: Parkinson’s disease
VAMP2: vesicle-associated membrane protein 2
vGLUT1: vesicular glutamate transporter 1

## Declarations

### Ethics approval

All animal protocols were approved by the Institutional Animal Care and Use Committee at the University of Alabama at Birmingham.

### Consent for publication

All authors read and approved the final manuscript for publication.

### Availability of Data and Materials

The datasets used and/or analyzed during the current study are available from the corresponding author on reasonable request.

### Author Contributions

CFB and LVD designed experiments and overall hypothesis. LVD, CFB, BH, VS, NZG, KK and DS conducted experiments and the analysis, LVD and CFB interpreted the results. DS and ED provided ExM protocols and training. KK and AW provided PF-360 chow. LVD, CFB, BH and VS wrote the manuscript, NZG, KK, AW, DS and EB revised the manuscript.

## Acknowledgments

We also thank the staff of the UAB HRIF, particularly Dr. Alexa Mattheyses for the support and valuable input during high resolution imaging.

## Figure Legends

***Table 1:*** List of statistical tests and results for each figure.

